# PUM2 binds SARS-CoV-2 RNA and PUM1 mildly reduces viral RNA levels, but neither protein affects progeny virus production

**DOI:** 10.1101/2025.06.23.661172

**Authors:** Nhi Phan, Yelizaveta Zaytseva, Chia-Ching Lin, Mitali Mishra, Weina Sun, Paulina Pawlica

**Author notes:** Department of Microbiology, New York University Grossman School of Medicine, New York, NY 10016. Contributed equally (Nhi Phan, Yelizaveta Zaytseva).

## Abstract

Pumilio proteins (PUM1 and PUM2) are essential post-transcriptional regulators of gene expression found across plants, animals, and yeast. They bind Pumilio Response Elements (PREs) on messenger RNAs (mRNAs) to modulate mRNA stability and translation. PUMs have been implicated in diverse cellular processes, including stem cell maintenance, neurogenesis, and cell cycle regulation. They have also been reported to negatively regulate innate immunity genes and to participate in viral RNA sensing. Previous high-throughput interactome studies revealed that PUMs bind SARS-CoV-2 RNA. We found that SARS-CoV-2 transcripts contain multiple conserved PREs, some of which are preferentially bound by PUM2. Surprisingly, altering PUM levels does not affect the production of progeny virions. However, depletion of PUM1 slightly increases intracellular viral RNA levels, suggesting that PUM1 either plays a mild antiviral role against SARS-CoV-2 or regulates a host factor that promotes viral replication. Notably, PUM1 also negatively regulates innate immunity gene expression both at steady state and during SARS-CoV-2 infection. Our findings support a complex immunomodulatory role for PUM1, acting both as a negative regulator of innate immunity genes and a mild inhibitor of SARS-CoV-2 RNA accumulation. However, in cell culture, these roles appear negligible based on viral progeny output. Whether the multiple PREs found in the SARS-CoV-2 genome contribute to evasion of PUM1 activity remains an open question.

## Introduction

Pumilio 1 (PUM1) and Pumilio 2 (PUM2) are conserved, sequence-specific RNA-binding proteins that function as essential post-transcriptional regulators of gene expression, playing critical roles in growth, development, neurological processes, and fertility^1, 2^. Mutations in these proteins have been linked to human diseases, including ataxia and various cancers^1, 2^. PUMs bind specific sequences known as Pumilio Response Elements (PREs; consensus 5′-UGUAHAUA, where H = A, C, or U) on messenger RNAs (mRNAs) via their highly conserved C-terminal homology domain (HD)^3^ to control mRNA stability and translation. Both PUM1 and PUM2 recognize the same PREs and largely share overlapping mRNA targets. However, PUM1 contains an additional N-terminal domain not present in PUM2, suggesting functional differences. Indeed, several studies have shown that the roles of PUM1 and PUM2 are not completely redundant^4-9^.

In addition to their role in mRNA regulation, PUMs participate in stress responses, as evidenced by their translocation into stress granules under various stress conditions, including viral infections^10, 11^. During infection, the innate immune system relies on pattern recognition receptors (PRRs), which detect pathogen-associated molecular patterns (PAMPs) and trigger immune signaling. Retinoic acid-inducible gene I (RIG-I)-like receptors (RLRs), including RIG-I and melanoma differentiation-associated protein 5 (MDA5) sense foreign RNA and initiate a signaling cascade that induces interferons, interferon-stimulated genes (ISGs), and pro-inflammatory cytokines^12^.

Two studies examined the roles of PUM proteins in regulating innate immunity and viral replication, reporting opposite, likely virus-specific, effects on viral replication^8, 10^. In 2014, Narita *et al*. reported that both PUMs act as positive regulators of RIG-I signaling by directly interacting with laboratory of genetics and physiology 2 (LGP2), another RLR, and enhancing its RNA-binding activity, thereby increasing the cell’s ability to detect foreign RNA^10^. The authors showed that depletion of PUMs increased, while their overexpression decreased, the RNA levels of Newcastle disease virus (NDV), a negative-sense paramyxovirus. Consequently, PUM depletion during NDV infection led to reduced induction of innate immunity genes such as interferon-beta (*IFN-β*) and C-X-C motif chemokine ligand 10 (*CXCL10*). Importantly, this function was independent of the PUM RNA-binding domain, suggesting that it did not involve canonical regulation of mRNA targets. In contrast, a 2017 study by Liu *et al*. found that PUM1, but not PUM2, acts as a negative regulator of innate immunity signaling by repressing expression of *LGP2, CXCL10*, and other immune genes^8^. This role was proposed to occur through post-transcriptional regulation of mRNA levels, especially that of *LGP2*. Notably, only about half of the genes upregulated upon PUM1 depletion were direct PUM1 targets, including *CXCL10* but not *LGP2*. Functionally, PUM1 depletion elevated basal expression of antiviral genes, which in turn suppressed replication of herpes simplex virus 1 (HSV-1), a double-stranded DNA virus. Thus, although both studies implicate PUMs in antiviral responses, they propose distinct mechanisms: one suggests a positive role for both PUMs in enhancing RIG-I signaling and antiviral activity, while the other describes PUM1 as a negative regulator of the antiviral state, implying a pro-viral role. These models are not mutually exclusive, as they employ different mechanisms; however, they indicate that the roles of PUMs in antiviral responses remain poorly understood and appear to be virus-specific.

In this study, we discovered that the SARS-CoV-2 genome harbors conserved PREs and demonstrated that some of these elements are preferentially bound by PUM2. We confirmed that PUM1 is mildly antiviral against NDV. In the context of SARS-CoV-2 infection, we showed that PUM1 depletion slightly increases viral RNA levels but does not affect viral progeny output. We also observed that loss of PUM1 leads to increased expression of innate immunity genes, including IFN-β, IFIT2, and the known PUM target CXCL10. These findings suggest that PUM1 has a complex immunomodulatory role; however, at least in cell culture, SARS-CoV-2 neither exploits PUMs to enhance replication nor is affected by PUM1’s antiviral or immunomodulatory functions.

## Results

### PUMs bind conserved PREs on SARS-CoV-2 transcripts

While analyzing the SARS-CoV-2 genome, we noticed that it contains multiple Pumilio Response Elements (PREs), with 6 located on the sense (genomic) strand and 4 on the negative-sense strand (**Fig. 1A**). Using GISAID’s^13^ nucleotide mutation data for SARS-CoV-2, we calculated genetic diversity scores for each 8-nucleotide (8-nt) sequence across the whole SARS-CoV-2 genome. Although SARS-CoV-2 genome is generally stable, our analysis revealed that the PREs are highly conserved, with the exception of 1 located on the antisense strand that exhibits elevated genetic variability (**Fig. 1B**). Given that antisense transcripts are less abundant and likely confined to replication compartments, we focused primarily on PREs located on the sense strand. To determine whether these viral PREs are bound by PUM proteins, we performed RNA immunoprecipitation (RIP) from lysates of SARS-CoV-2-infected Calu-3 cells. Interestingly, 4 PREs located on the sense strand were found to be preferentially bound by PUM2, as compared to the negative control, the *N* transcript, which lacks PREs (**Fig. 1C**). SARS-CoV-2 transcripts showed lower enrichment than the known host PUM target cyclin-dependent kinase inhibitor 1B (*CDKN1B*)^9, 14^ or the long non-coding RNA *NORAD*, which sequesters PUMs via ∼20 PREs^15^. Some of the predicted PRE-containing viral transcripts were not bound by PUMs, which may be due to occlusion of binding sites by RNA structure or other RNA-binding proteins. Importantly, three independent high-throughput studies investigating protein interactions with SARS-CoV-2 transcripts identified PUM proteins as associated with viral RNA. Two of these studies detected PUM2^16, 17^, and all three identified PUM1^16-18^. Since both PUM1 and PUM2 recognize the same RNA motif (PRE), it is likely that PUM1 also binds SARS-CoV-2 RNA, even though we primarily detected PUM2. In summary, our data and those of others show that SARS-CoV-2 harbors conserved PREs, some of which are bound by PUM proteins, particularly PUM2.

**Figure 1.**
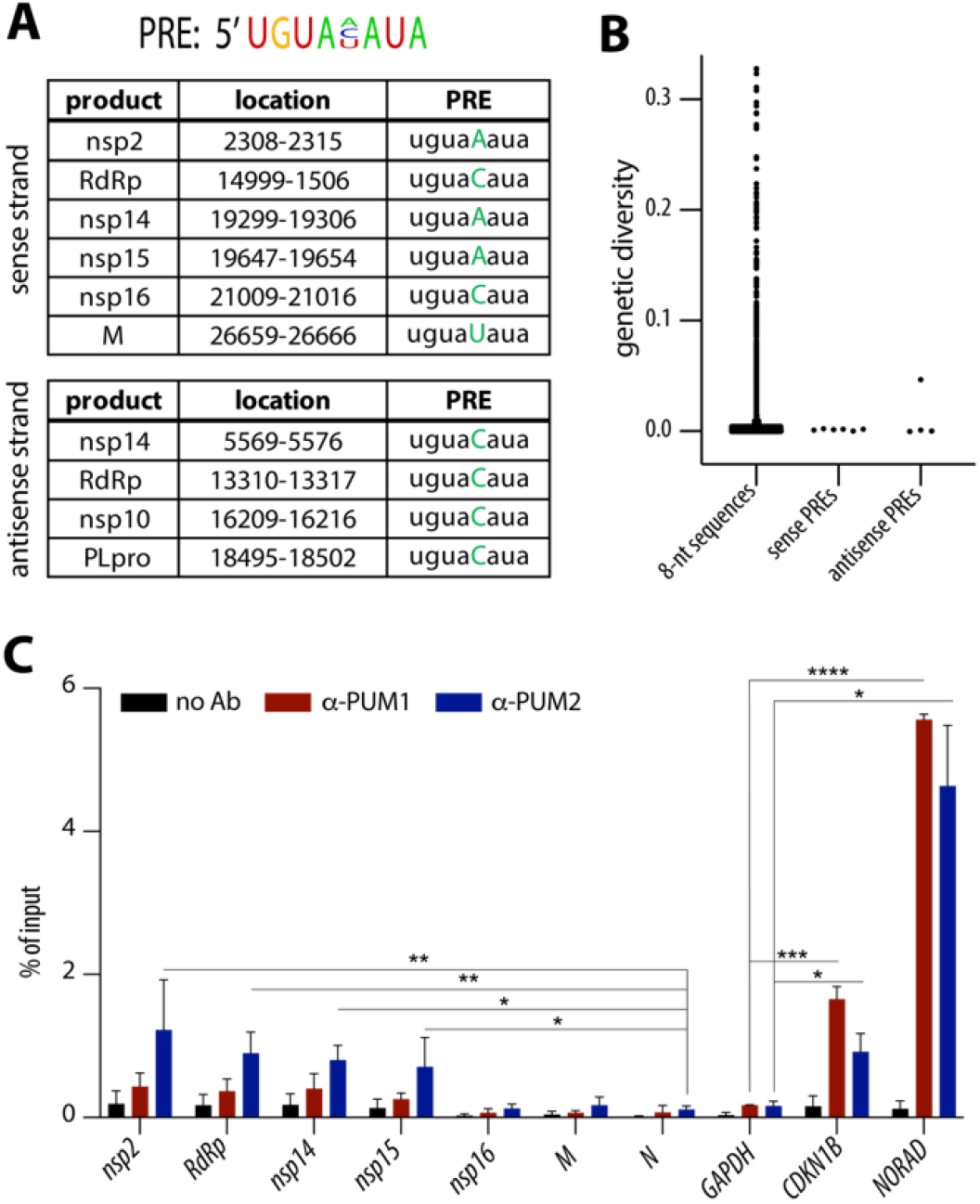
PUM2 binds conserved PREs on SARS-CoV-2 transcripts. **(A)** Top: Consensus motif of the Pumilio response element (PRE). Below: List of all PREs and their location in the SARS-CoV-2 genome. **(B)** Most of the PREs within the SARS-CoV-2 genome are genetically stable. Mutation rates for 8-nt sequences across the genome were calculated using diversity statistics from GISAID data. **(C)** SARS-CoV-2 transcripts are preferentially bound by PUM2. The graph shows mean values from RNA immunoprecipitations of PUM1 and PUM2 from Calu-3 cells infected with SARS-CoV-2 WA1 for 48 h at MOI 0.05 (n = 3; error bars indicate SD). Significance was calculated using a ratio paired t-test by comparing values to the negative control: *GAPDH* for the host transcripts and *N* for the viral RNAs. * indicates a p-value of < 0.05, **p < 0.01, ***p < 0.001, and ****p < 0.0001. *CDKN1B* is a known PUM target, while *NORAD* contains ∼20 PREs.

## PUM1 depletion mildly increases SARS-CoV-2 RNA levels but does not affect progeny virus production

Since PUM proteins have been shown to modulate innate immunity, and both our data and previous studies have detected their binding to the viral genome, we reasoned that the virus may either exploit PUMs to promote its replication or sequester them to inhibit their antiviral function. To assess the impact of PUM depletion on SARS-CoV-2 replication, we used CRISPR-Cas9 to generate single PUM knockouts in A549-hACE2 cells, with two independent clones for each protein. Because cells with double PUM knockout exhibit growth delays due to prolonged de-repression of PUM targets^16, 17^, we employed a strategy in which one PUM was knocked out and the other was depleted using siRNAs (**Fig. 2A**). We measured cell viability via DNA synthesis and confirmed that cells continued to proliferate despite transient double PUM depletion (**Fig. 2B**). We further validated our system by confirming that depletion of both PUMs leads to de-repression of their known targets: *CDKN1B* and Tousled-like kinase 1 (*TLK1*)^9, 14, 19^ (**Fig. 2C**). Since Narita *et al*. reported that PUMs have antiviral activity against NDV^10^, we asked whether this observation is reproducible in A549 cells. We confirmed that PUM1, but not PUM2, negatively regulates NDV replication, as measured by viral titers (**Fig. 2D**). In the aforementioned study, NDV replication was measured by viral RNA levels—a more sensitive method capable of detecting subtle effects, such as those potentially mediated by PUM2. Surprisingly, in the context of SARS-CoV-2 infection, neither single nor double PUM depletion affected viral replication, as measured by progeny virion production (**Fig. 2E**). Interestingly, viral RNA levels were slightly upregulated in PUM1-depleted cells, but not in PUM2-depleted cells, suggesting that PUM1 may have a mild antiviral role against SARS-CoV-2 or may regulate a host factor that promotes viral replication (**Fig. 2F**). If PUM1 is indeed antiviral, the lack of major activity against SARS-CoV-2 might indicate that the virus has evolved a strategy to evade this function. In summary, we confirm a mild antiviral activity of PUM1 against NDV, and observe a subtle inhibitory effect on SARS-CoV-2 RNA levels; however, PUM depletion has no significant impact on progeny virus production.

**Figure 2.**
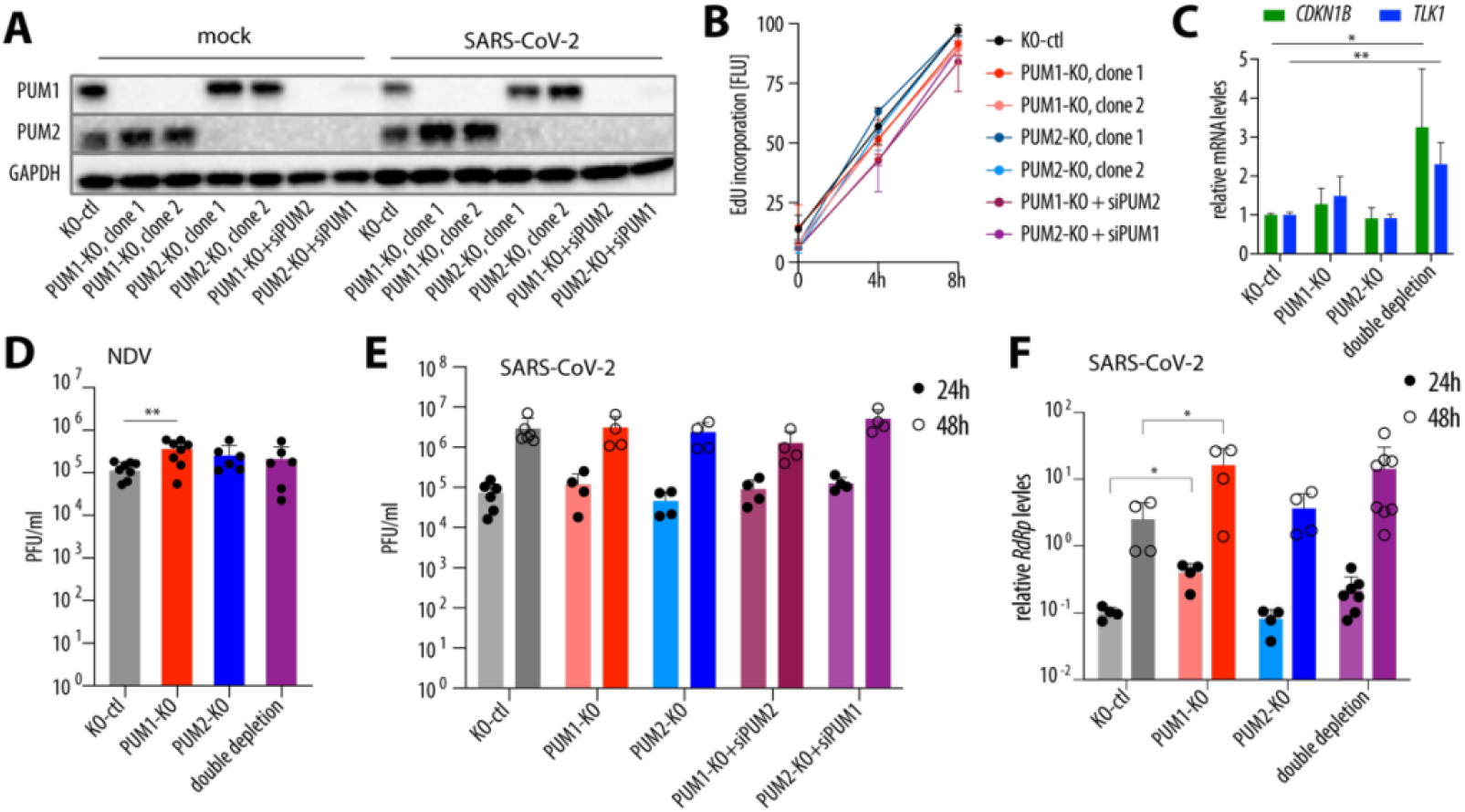
SARS-CoV-2 replication is largely unaffected by PUM depletion. **(A)** Validation of successful PUM depletion. A representative Western blot from uninfected and SARS-CoV-2-infected cells at 48 hpi. Clonal cell lines derived from A549-hACE2, either control (KO-ctl) or single knockouts (KO) of PUM1 or PUM2, were treated with siRNAs (siPUM1, siPUM2, or non-targeting control). 48h post siRNA treatment, cells were either infected or mock-infected with SARS-CoV-2 WA1 at an MOI of 0.5. **(B)** Cells with PUM depletion are viable. Viability was assessed by EdU incorporation into newly synthesized DNA. Data represent the average of two experiments performed in three biological replicates. **(C)** PUM double depletion de-represses known PUM targets, *CDKN1B* and *TLK1*. *p < 0.05 and **p < 0.01 (ratio paired t-test, n =3). **(D)** PUM depletion modestly increases NDV replication. A549-hACE2 cells, either controls or with single or double PUM depletion, were infected with the NDV LaSota strain at MOI 0.1. Viral replication was measured by plaque assay at 24 hpi. ** – p < 0.01 (paired t-test, n = 6). **(E)** SARS-CoV-2 progeny virus production is unaffected by PUM depletion. A549-hACE2 cells, either controls or with single or double PUM depletion, were infected with SARS-CoV-2 WA1 at an MOI of 0.5, and replication was measured by plaque assay. Graphs represent results from at least two experiments, each including two control cell lines, two clonal knockouts of each PUM, and four double depletion conditions. **(F)** SARS-CoV-2 RNA levels are slightly increased by PUM1 depletion. The experiment was set up as in **(E)**, and intracellular viral RNA levels were measured by RT-qPCR and normalized to *GAPDH*. *p < 0.05 (paired t-test). Error bars indicate SD. hpi – hours post-infection; EdU – 5-Ethynyl-2′-deoxyuridine; PFU – plaque-forming units; RdRp – RNA-dependent RNA polymerase.

### PUM overexpression does not affect SARS-CoV-2 progeny virus production

We also investigated whether PUM overexpression affects SARS-CoV-2 replication. A549 cells are notoriously difficult to transfect, and we observed that stable transduction leads to rapid loss of exogenous PUM expression, possibly due to silencing by the HUSH complex^20^. Additionally, PUM protein levels are tightly regulated through mechanisms including auto- and cross-regulation via PREs within their own mRNAs^1, 21^, as well as sequestration by the long non-coding RNA *NORAD*^22-24^. Thus, to assess the effect of PUM overexpression, we transiently transduced A549-hACE2 cells with lentiviral vectors carrying PUM constructs (**Fig. 3A, B**). We first confirmed successful overexpression by demonstrating downregulation of known PUM targets, *CDKN1B* and *TLK1*. In the context of SARS-CoV-2 infection, PUM overexpression did not substantially affect viral titers (**Fig. 3C**). In summary, our data show that overexpression of PUM proteins plays a negligible role in production of SARS-CoV-2 progeny virions.

**Figure 3.**
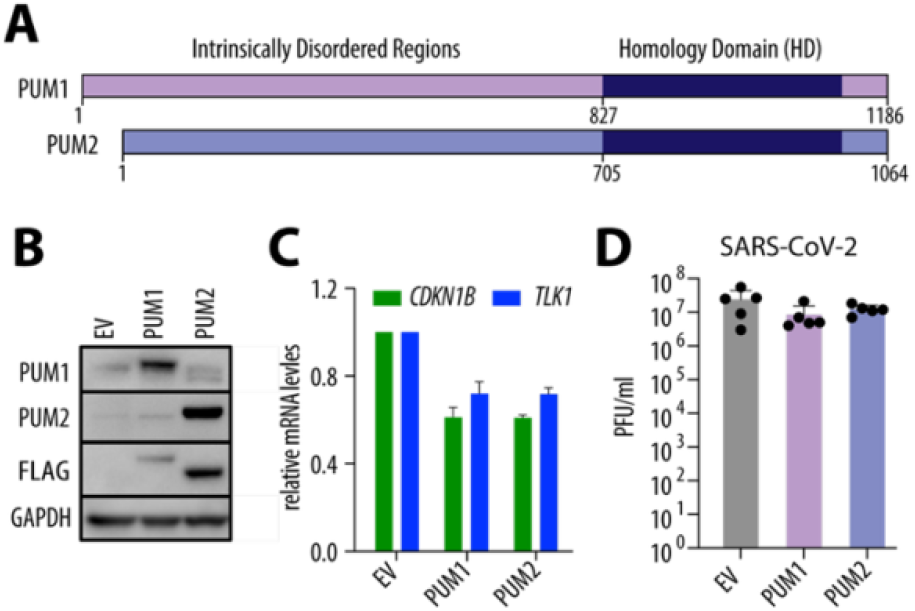
PUM overexpression does not affect SARS-CoV-2 replication. **(A)** Schematic representation of PUM proteins. **(B)** Representative Western blot from A549-hACE2 cells transiently transduced with the indicated FLAG-tagged constructs. **(C)** PUM overexpression represses known PUM targets, *CDKN1B* and *TLK1* (n = 2). **(D)** PUM overexpression does not impact SARS-CoV-2 replication. A549-hACE2 cells were transduced with the indicated constructs and infected with SARS-CoV-2 WA1 at MOI 0.5 for 48 h. Viral replication was measured by plaque assay (n = 5). Error bars represent SD. hpi – hours post-infection; PFU – plaque-forming units.

### PUM1 negatively regulates innate immunity genes, but this activity is not enhanced by type I IFN treatment

Two previous studies reported that PUM proteins regulate innate immunity genes during NDV and HSV-1 infections; however, the directionality of this regulation differed between the studies^8, 10^. We thus investigated how PUM depletion affects the levels of *CXCL10, IFN-β, ISG15*, and *IFIT2* at steady state and during SARS-CoV-2 infection (**Fig. 4A**). At steady-state levels (mock), the absence of PUM1 led to higher levels of most tested genes, with the exception of *ISG15*; however, statistical significance was reached only for *IFIT2*. During SARS-CoV-2 infection, induction of the antiviral program was inhibited by the virus, consistent with its known ability to suppress IFN signaling^25-29^. There was a general trend that, in the absence of PUM1, innate immunity genes were expressed at higher levels compared to control cells. The genes that reached statistical significance at any time point during infection were *IFN-β* and *CXCL10*. This is in agreement with findings by Liu *et al*., who proposed that PUM1 acts as a negative regulator of innate immunity genes^8^. We then tested whether PUM1 depletion alters the antiviral state induced by 24 h of IFNα treatment and found no change. (**Fig. 4B**). Consistently, the inhibitory effect of IFNα pretreatment against SARS-CoV-2^30, 31^ was unaltered by PUM1 depletion. Overall, these results suggest that although PUM1 negatively regulates the expression of innate immunity genes, these effects are negligible for SARS-CoV-2 replication in cell culture.

**Figure 4.**
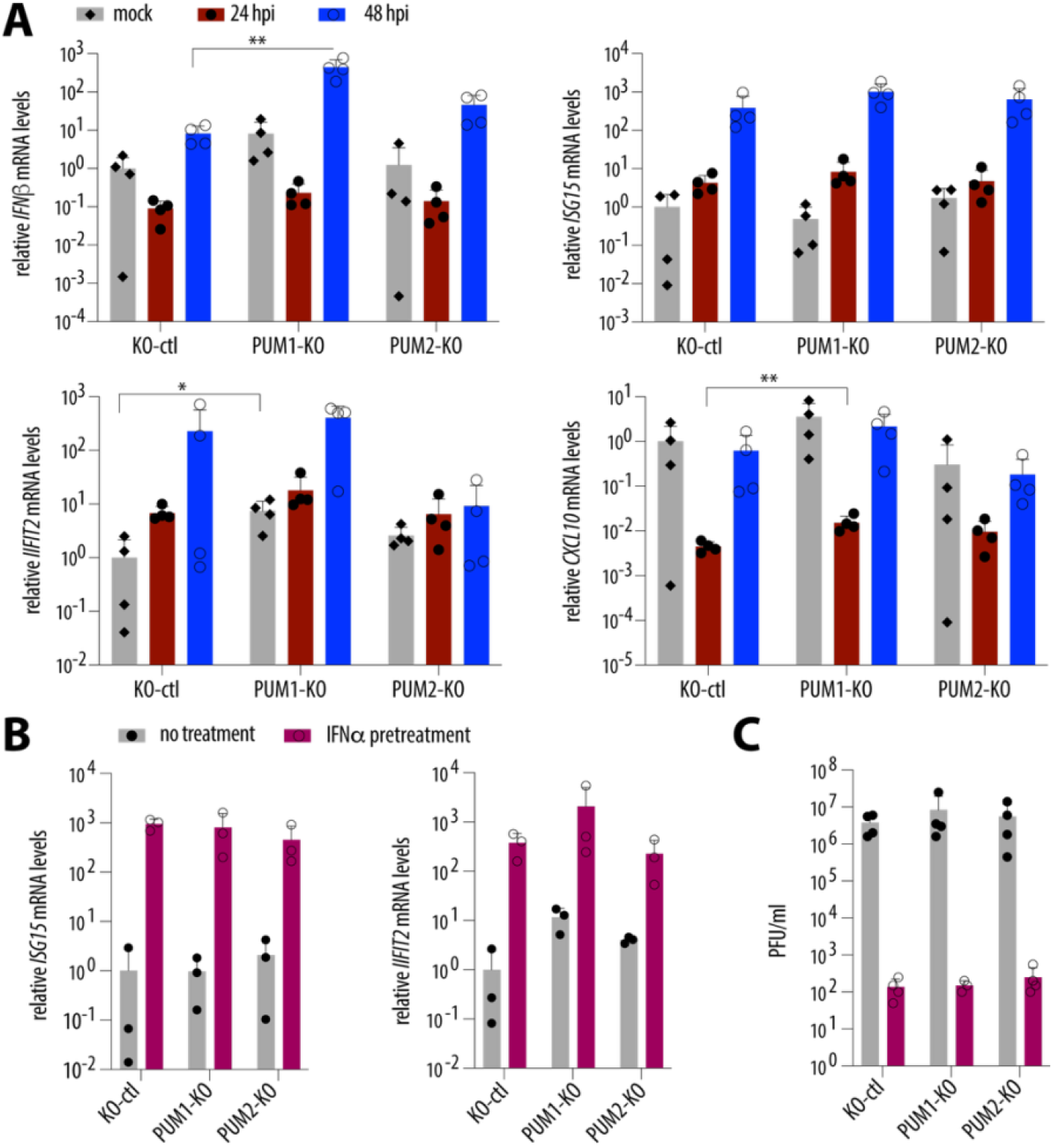
PUM1 negatively regulates innate immunity genes, but this activity is not enhanced by type I IFN treatment. **(A)** PUM1 depletion negatively regulates expression of innate immunity genes during steady state and SARS-CoV-2 infection. A549-hACE2 cells, either control or with single PUM depletions, were infected with SARS-CoV-2 WA1 at an MOI of 0.5, and mRNA levels of *CXCL10, IFN-β, ISG15*, and *IFIT2* were measured by RT-qPCR. Results were normalized to *GAPDH* and then to mock. *p < 0.05, **p < 0.01, (paired t-test, n = 4). **(B)** PUM1 depletion does not enhance the antiviral state induced by IFNα treatment. A549-hACE2 cells, either control or with single PUM depletions, were treated with 103 U/ml of IFNα for 24 h, after which RT-qPCR was performed (n = 3). **(C)** Single PUM depletion does not affect the impact of IFNα pretreatment on SARS-CoV-2 replication. A549-hACE2 cells, either control or with single PUM depletions, were pre-treated overnight with 103 U/ml of IFNα and infected at an MOI of 0.5 with SARS-CoV-2 WA1. Viral replication was assayed 48 h post-infection. Error bars represent SD. hpi – hours post-infection; RdRp – RNA-dependent RNA polymerase.

## Discussion

In this study, we identified genetically stable PREs present in the SARS-CoV-2 genome and showed that some of them are preferentially bound by PUM2 (**Fig. 1C**). Interestingly, other high-throughput studies have identified both PUM proteins being associated with SARS-CoV-2 transcripts^16-18^. PUM1 and PUM2 are known to interact with each other ^19^, which may explain why the presence of one protein is often accompanied by the other.. Also, since both PUM1 and PUM2 recognize the same RNA motif (PRE), it is likely that both bind to SARS-CoV-2 RNA, although one may exhibit a preference depending on the surrounding RNA context. On the other hand, the selective binding of PUM2 might be biologically relevant, as in our experiments this bias is unlikely to result from technical issues such as antibody quality—both anti-PUM1 and anti-PUM2 antibodies efficiently pulled down host transcripts, with anti-PUM1 performing even better. (**Fig. 1C**). One hypothesis for the preferential binding of PUM2 is that it may serve to shield the viral genome from PUM1’s antiviral properties.

Here, we confirmed that PUM1 is modestly antiviral against NDV (**Fig. 2D**). Although we did not observe major antiviral activity of PUM1 against SARS-CoV-2, as measured by progeny virion production, we detected slightly higher levels of intracellular viral RNA in the absence of PUM1, which could suggest the presence of residual antiviral function (**Fig. 2E, F**). It is unclear whether the mechanism underlying these antiviral properties is the same as for NDV, since Narita *et al*.^*10*^ proposed that PUMs participate in foreign RNA sensing and thereby enhance downstream signaling. In contrast, we observed that PUM1 negatively regulates innate immunity, as proposed by Liu *et al*. ^8^ (**Fig. 4A**). Thus, our results are partially consistent with the two previously published studies and suggest that PUM1 may have at least two distinct functions: one antiviral and one involving negative regulation of innate immunity genes. It is also possible that PUM1 is not directly antiviral, but instead regulates the expression of a host factor that promotes viral replication. However, in the context of SARS-CoV-2, regulatory roles of PUM1 appear to be negligible, at least in cell culture. A limitation of this study is that all experiments were conducted in cell culture, and it remains unclear whether PUM1 plays a significant role in viral infection within an organism.

Our study expands the understanding of the immunomodulatory roles of PUM1 in the context of viral infection, specifically during SARS-CoV-2 replication. There is an intriguing discrepancy regarding the role of PUM1 in viral infections: it appears antiviral against NDV, proviral for HSV-1, and largely neutral for SARS-CoV-2. Two possible explanations may account for this. First, these viruses belong to distinct families and produce different PAMPs, which are recognized by different PRRs. This could influence PUM1’s involvement in foreign RNA sensing, as proposed by Narita *et al*.^10^. Second, given the antiviral function of PUM1 against NDV—a virus whose genome lacks PREs—it is tempting to speculate that SARS-CoV-2 and HSV-1, both of which contain multiple PREs, may use these elements to evade PUM1’s antiviral function. Mechanistically, PRE-mediated evasion could occur through preferential binding of PUM2, shielding the viral genome from PUM1’s effects, or through sequestration of PUM1, preventing its interaction with LGP2 and impairing its role in RNA sensing. These hypotheses, however, are difficult to test, as they would require introducing multiple mutations into viral genomes. At present, it remains unclear whether PUM1 has major antiviral activity against viruses, as even its activity against NDV appears modest at best.

## Author Contributions

P.P. designed the study. P.P., N.P., C.-C.L., Y.Z., and M.M. performed the experiments. W.S. provided guidance on the NDV experiments. P.P. wrote the manuscript.

## Acknowledgements

We thank Adolfo García-Sastre for providing the A549-hACE2 cells, and Brad Rosenberg and Vincenzo Alessandro Gennarino for helpful discussions.

## Materials and Methods

### Cells

Calu-3 (ATCC) cells were cultured in EMEM (ATCC) and supplemented with 10% fetal bovine serum (FBS; HyClone), penicillin-streptomycin-glutamine (Pen/Strep/Glut; GIBCO), and 1 mM sodium pyruvate (GIBCO). A549-hACE2 (a kind gift from Adolfo García-Sastre) cells were cultured in DMEM (GIBCO) with 10% FBS, Pen/Strep/Glutamine, and 1 µg/ml blasticidin (GIBCO). HEK293T (ATCC) and Vero E6 (ATCC) cells were cultured in DMEM (GIBCO) with 10% FBS and Pen/Strep/Glut.

### Generation of PUM knockout cell lines

PUM-targeting sequences (see Table 1) were cloned into lentiCRISPRv2 (Addgene, #49535). Lentiviral vectors were prepared in HEK293Ts by transfecting guide-containing lentiCRISPRv2 together with helper plasmids pMDG.2 and psPAX2 (Addgene, #12259, #12260). A549-hACE2 cells were transduced with lentiviral vectors at low MOI followed by puromycin (GIBCO) treatment and clonal selection. Each clone was verified by Western blotting and Sanger sequencing of the gene loci. Two knockout clonal cell lines for each PUM were used.

**Table 1.**
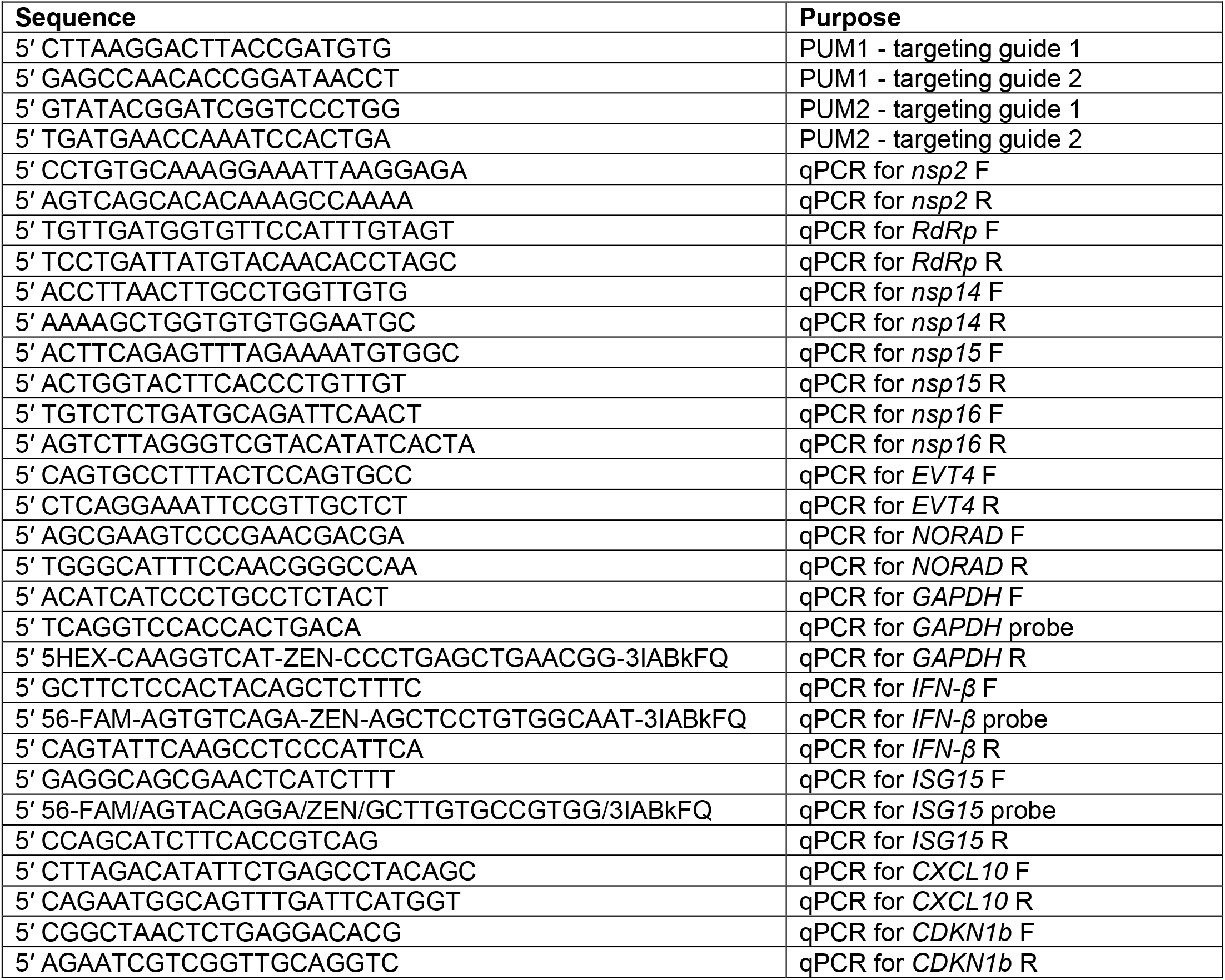
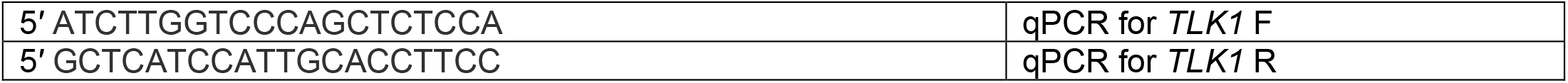

### PUM double depletion and overexpression

For PUM double depletion, clonal cell lines derived from A549-hACE2, either controls (KO-ctl, 2 separate cell lines) or single knockouts of either PUM1 (clone 1 or 2) or PUM2 (clone 1 or 2), were additionally transfected with siRNAs (siCtrl, siPUM1, or siPUM2) for 48 h and then infected. Specifically, 2×105 cells were seeded in a well of a 6-well plate and transfected next day using RNAiMAX (Invitrogen) with a 10 nM mix of 2 siRNA per gene (Silencer RNA, Ambion, # s18681, s18680, s23671, s23673). For PUM overexpression, PUM reference sequences were cloned into pLVX-Puro (Takara) between the NotI and BamHI sites. Empty pLVX control or pLVX containing PUM constructs were transfected into HEK293Ts together with helper plasmids to generate lentiviral vectors. A549-hACE2 cells were seeded at 2×105, transduced after 24 h with 1 ml of lentivirus-containing media, and then infected after additional 48 h. Successful depletion and overexpression were validated for each experiment.

### SARS-CoV-2 stocks and infections

SARS-CoV-2 stocks of isolate USA-WA1/2020 (BEI) were generated from passage 1 on VERO-E6 cells. Virus titers were determined by plaque assay as described previously^32^. A549-hACE2 cells were infected at MOI of 0.5, and Calu-3 cells were infected at an MOI of 0.1. For IFNα (PBL assay science) pretreatment, cells were incubated overnight with 103 U/mL and infected the next day. All infectious growth was performed in a Biosafety Level 3 laboratory and approved by the Mount Sinai Biosafety Committee.

### NDV stocks and infections

NDV stocks LaSota strain were generated and titrated as previously described^33,34^. Briefly, recombinant NDV was rescued using plasmids pNDV-LaSota, pTM1-NP, pTM1-P, pTM1-L, and pCAGGs-T7opt in BSRT7 cells for 48 h. Afterward, the homogenized cells and supernatant were injected into 9-to 11-day-old SPF embryonated chicken eggs (AvsBio), which were incubated at 37°C for 2–3 days. Viruses in the allantoic fluids were purified through ultracentrifugation on a 20% sucrose cushion and titrated on Vero E6 cells. For infection, an MOI of 0.1 was used. Viruses were diluted in PBS containing 0.2 % BSA (MP Biomedicals), Pen/Strep, calcium chloride, and magnesium chloride. After a 1-h incubation, the inoculum was removed, and DMEM containing 0.3% BSA, Pen/Strep, and 0.2 μg/ml tosylsulfonyl phenylalanyl chloromethyl ketone (TPCK) trypsin was added. 24 hpi, the supernatants were harvested, and viral titers were measured by plaque assay on Vero E6 cells.

### RNA Immunoprecipitation (RIP)

RIP was performed as previously described^32^. Briefly, 5×103 Calu-3 cells were infected with SARS-CoV-2 WA1 at MOI of 0.05 for 48 h. Cells were lysed in 1 ml of IP Lysis Buffer (Pierce) in the presence of RNase inhibitor (NEB) and Protease Inhibitor Cocktail (Pierce). Lysates containing 400 µg of protein were incubated with 30 µl of Dynabeads Protein G (Invitrogen) coupled to 10 µg of antibody (anti-PUM1 or anti-PUM2, Bethyl), or with beads only as a no-antibody control. After IP, the beads were washed twice in Pierce Lysis Buffer and then 5 times in NET-2 buffer (50 mM Tris [pH 7.5], 150 mM NaCl, and 0.05% Nonidet P-40). Beads were resuspended in TRIzol (Invitrogen), and RNA was extracted and analyzed by RT-qPCR.

### RT-qPCR

Total RNA was extracted from TRIzol according to the manufacturer’s recommendations. 2 µg of RNA was subjected to DNase I digestion (Promega) and purified with acid phenol. 1 µg of DNase -treated RNA was reverse transcribed using ProtoScript II and random hexamers. qPCR was performed using either SYBR Green qPCR Master Mix (Thermo Fisher) or Luna qPCR Master Mix (NEB). The reactions were performed according to the manufacturer’s instructions. Sequences are listed in Table 1.

### Edu incorporation

The cell proliferation assay was performed using the Click-iT EdU Proliferation Assay (Invitrogen) according to the manufacturer’s instructions. Briefly, 10^4^ cells from each PUM knockout clone, pretreated with siRNAs for 48 hours, were plated in a 96-well plate. The following day, the cells were incubated with 10 µM EdU for the indicated times. Afterward, the cells were fixed, labeled with the Click-iT reagent, washed, blocked with 1.5% BSA, and incubated with Amplex UltraRed for 15 minutes. The signal was measured using a Synergy H1m BioTek instrument.

### Western blot analysis

Cells were lysed in IP Lysis buffer (Pierce) containing protease inhibitor cocktail (Pierce). ∼20 µg of total protein was separated on a gradient SDS-PAGE gel (Invitrogen), and electrotransferred to a PVDF membrane (BioRad). After blocking with 5% milk in TBST (20 mM Tris [pH 7.5], 150 mM NaCl, 0.1% Tween 20), the membrane was probed with the appropriate antibodies and detected using ECL Plus Western Blotting Substrate (Pierce) on a ChemiDoc MP Imaging System. Primary antibodies used were anti-FLAG M2 (1:1000, F1804, Sigma-Aldrich), anti-PUM1 (1:2000, ab92545, Abcam or A300-201A, Bethyl), anti-PUM2 (1:1000, A300-202A, Bethyl, or ab92390, Abcam) and anti-GAPDH (1:5000, G9545, Sigma-Aldrich). Secondary antibodies used included Sheep Anti-Rabbit IgG Antibody, HRP conjugate (1:5000, AP510P, Sigma Aldrich) and Rabbit Anti-Mouse IgG Antibody, HRP conjugate (1:5000, AP160p, Sigma-Alrich).

### Bioinformatic analysis

All-time per-nucleotide diversity statistics for SARS-CoV-2 (NC_045512.2) were downloaded from GISAID^13^ on July 9, 2024. Missing values were set to zero. Rolling averages of diversity were calculated across 8-nt windows using a 1-nt step to smooth local variation, and the resulting values were Z-score normalized to identify regions of unusually high or low conservation. Diversity scores for PRE elements in the SARS-CoV-2 genome were then compared to those of all other 8-nt windows.

